# Role of Efferocytosis in Oral Lichen Planus

**DOI:** 10.1101/2021.08.09.455749

**Authors:** Thavarajah Rooban, Kannan Ranganathan

## Abstract

**Background:** Oral lichen planus (OLP) is a chronic, immune mediated interface mucositis of oral mucosa. Though the apoptosis of keratinocytes is a feature of OLP, not much is known about the clearance of cell debris (efferocytosis) resulting from apoptosis. We postulate that there is a defective or delayed efferocytosis in OLP, which may have a role in modulating the immune response in OLP.

**Methods:** Published mRNA expression of tissue of 14 patients with OLP and 14 cases of normal tissues were subjected to differential analysis (DE) and a list of DE genes identified. From this list, the genes that involved in efferocytosis were collated, compared and their interactions are typed.

**Result:** In all, two studies fulfilled the inclusion and exclusion criteria. On combining the data, 1486 genes were significantly different between OLP and normal tissues. 28 of these 1486 genes are were associated with efferocytosis of which the suppression of LRP1, LDLR, ANAX2, C2, PBX1, PDCD4, S1PR5, CX3CL1, STAT6 and Wnt3A is indicative of defective or delayed efferocytosis in OLP. The role of pathways and associations were analyzed and is presented here.

**Discussion and Conclusion:** The study revealed that certain key genes mRNAs that are associated with efferocytosis are altered in OLP. They could delay or lead to defective efferocytosis. Studying such genes in detail could provide deeper understanding of the pathogenesis of the disease and the discovery of therapeutic targets.

## BACKGROUND

Oral lichen planus (OLP) is a non-infectious, T-cell–mediated, chronic inflammatory disorder of oral mucosa and lamina propria. Systemic conditions such as psychological stress, anxiety, hepatitis C virus infection, hypertension, diabetes, graft vs host disease, and thyroid dysfunction have been associated with OLP.^[1-3]^ It is a part of spectrum of cell-mediated immunopathogenesis or a dysregulated T cell-mediated disorder to exogenous trigger or a dysregulated immune response to autologous keratinocyte antigens (autoimmune).^[1-3]^ OLP is associated with both CD4+ (“helper”) and CD8+ (“cytotoxic”) T cells. The CD4+T cells produce Th1 factors while the CD8+ cytotoxic T cells kills the host keratinocytes. CD8+ T cells are observed in association with epithelial-connective tissue interface and adjacent to apoptotic keratinocytes.^[1-3]^ This aggregation of inflammatory cells near the mucosal interface is hall mark of the OLP.^[4-8]^ Apoptosis and necroptosis occur in lichen planus.^[4,5]^ Destruction of keratinocytes, in OLP is a culmination of auto-reactivity and/or as a by-stander effect.^[4,5]^

Dead human cells, if not properly cleared, trigger endosomal receptors such as TLR7,-8, and -9 resulting in downstream activation of several signaling pathways leading to production of pro-inflammatory cytokines.^[9,10]^ In OLP, like several autoimmune disorders, the levels of circulating pro-inflammatory cytokines are elevated.^[6]^

The apoptotic cells are cleared by phagocytes, a process termed ‘efferocytosis’. Efferocytosis happen without any inflammatory or immune reactions. This scavenging process is vital for the upkeep of normal tissue homeostasis. Several chronic inflammatory diseases are associated with defective efferocytosis. ^[9]^

Under certain conditions, apoptosis, especially in immunological challenges, can be similar to Immune-mediated cell death(ICD) too.^[9]^ Cell death is related to the activation of the damage-associated molecular patterns (DAMPs) or pattern-associated molecular patterns (PAMPs), which are a result of pre-mortem (just before death) stress response.^[10]^

Efferocytosis is different from simple phagocytosis and involves several phases such as apoptotic cell(AC) finding (“find-me”), binging, internalization (“eat-me”) and degradation.^[11]^ In the first phase, ACs attract immune cells which releases chemokines like CX3CL1 and lipids (like sphingosine 1-phosphate(S1P). In the second phase, phagocytes engage ACs through cell-surface receptors that either directly bind molecules on the surface(stabilin 1, stabilin 2, BAI1, TIM family, LRP1) or bind bridging molecules that interact with the AC surface (TYRO3, AXL and MER proto-oncogene tyrosine kinase -MERTK), the αvβ3 and αvβ5 integrins and the scavenger receptor CD36). The most important AC-associated signal is externalized phosphatidylserine (PS).^[11]^ Recognition of PS on ACs by PS receptors on phagocytes is the hall mark signal for efferocytosis. This receptor-ligand interactions are influenced by complements such as C1q, C2, CD91, LRP1 as well as other PS-binding proteins like ANXA5 and ANXA2^.[11-13]^

LRP1 is a type 1 trans-membrane protein that has several functions. It serves as a scavenger receptor (by binding and internalizing ligands for degradation and recycling), signaling receptor (interact with proteins and transduces intracellular signals involved in biological processes and regulating nuclear signaling) and as scaffold receptor (modulating membrane proteins such as integrins and receptor tyrosine kinases). It is known to suppress cell death as well as participate in efferocytosis. With ABCA1 and ABCG1, LRP1mediates efferocytosis which is known to activate PPARG, resulting in cholesterol efflux.^[14]^ LRP1 is pro-efferocytotic entity and its reduction may reduce the efferocytosis and prevents cholesterol efflux.^[11]^ Increased APOE promotes efferocytosis via the LRP1 complex through the PS. APOE binds to several receptors, such as LRP1 and LDLR. They can bind to ligands, signal downstream effects that leads to efferocytosis.^[11,14]^ Proteins and factors such as PLAUR, vitronectin, integrins, AKT1, IL1B, IL2RA, CASP1, CD163, IL10, PBX1, ADAM9, STAT6, Wnt-3A, PDCD4, S1PR5, KLK7, IDO1, ErbB2, BCL6 and NRP2 also can influence the efferocytosis.^[15-36]^ For adequate resolution of inflammation, efferocytosis is essential.^[11]^ [Figure-1, table-1]

**Figure-1:**
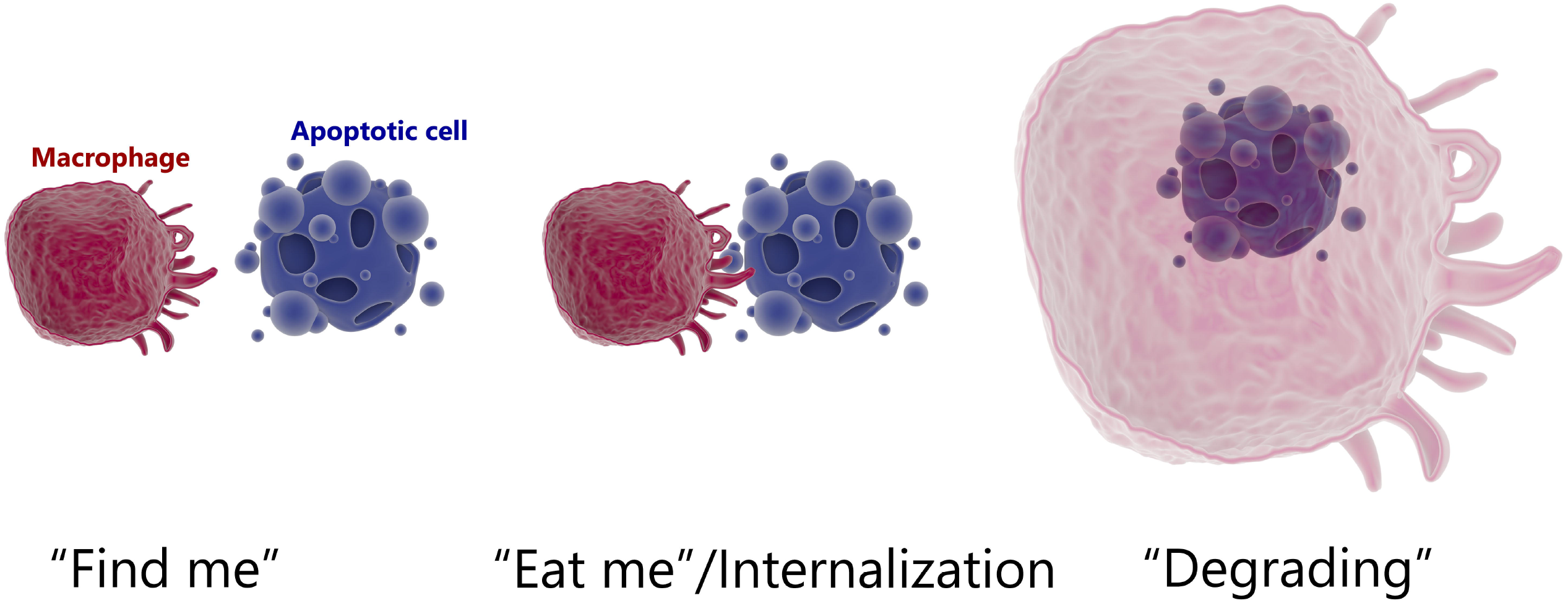
Schematic representation of Stages of efferocytosis

**Table-1:**
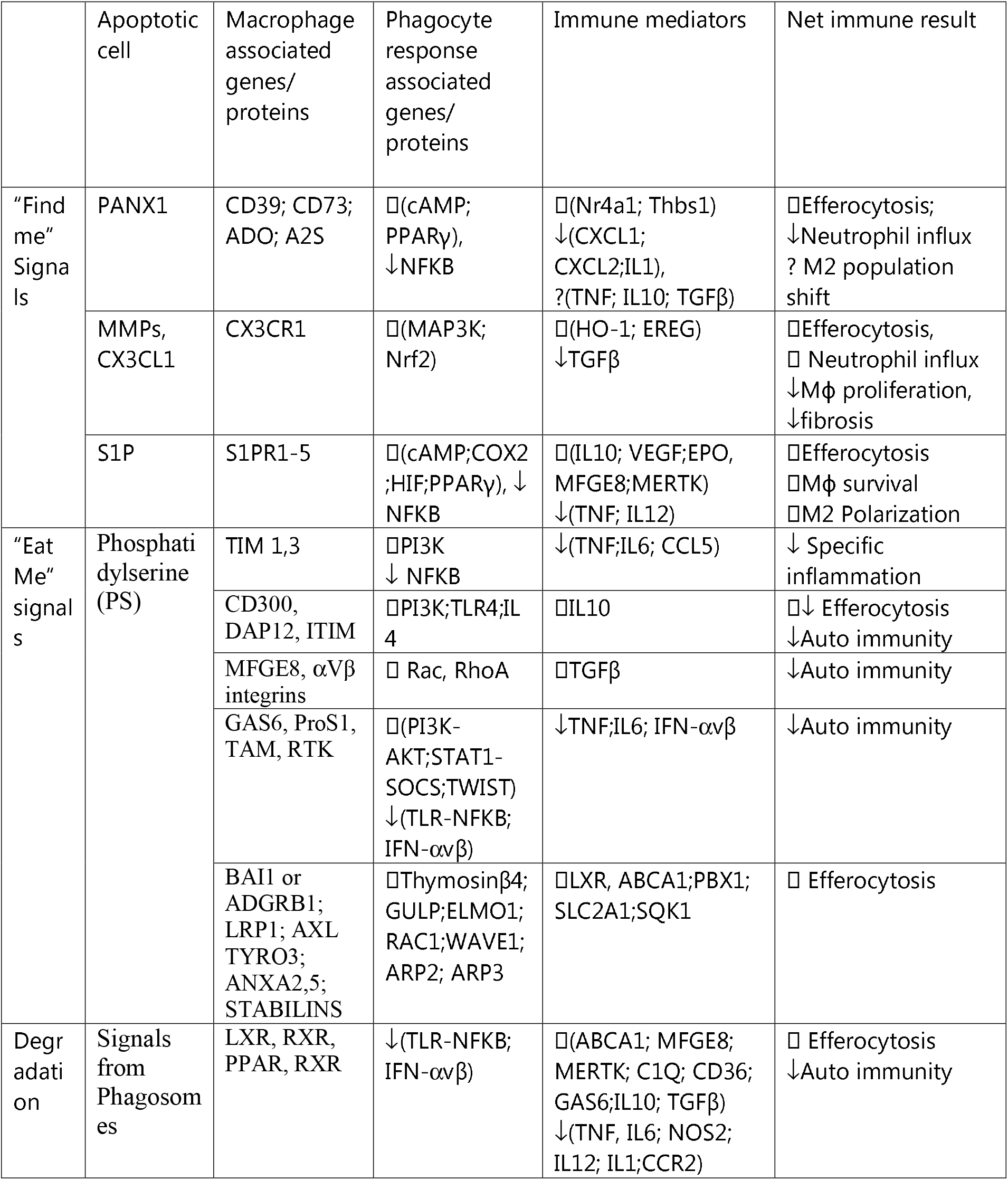
Interactions of genes/ proteins in efferocytosis

Though OLP has several features of autoimmune disorders, it has not been studied under the prism of efferocytosis process. This manuscript intends to explore the possibilities and spectrum of alteration in expression of mRNA of genes-proteins associated with efferocytosis process in OLP tissue as compared to normal mucosal tissue.

## MATERIAL AND METHODS

### Cell death and Cell debris removal related Gene Identification

Genes related to efferocytosis were enumerated from previously published work. The human gene ontology (http://geneontology.org) and www.genecards.org were used to collect genes associated with efferocytosis (GO:2000427, GO:0043277, GO:2000426, GO:2000425) along with literature mining using “Function Related Gene and Gene Network” option of Geneclip3.^[37]^ Data were mined only for human genes associated with efferocytosis, from Medline indexed journals published between 1980 to 2019.

### Data collection

Collection of the data for this study was carried in later half of January 2020 out by searching publicly available gene expression database – Gene Expression Omnibus (GEO), an International repository database repository of gene expression data and hybridization arrays, chips, microarrays hosted by the National Center for Biotechnology Information, USA. The search term used were “Lichen planus” with appropriate filter of organism (Homo sapiens), study type (expression profiling by array) and entry type (Dataset/Series). The entries were screened. Duplicates and irrelevant studies were excluded. Remaining studies were further refined using the inclusion criteria (below) to reach the 2 final studies included in present analysis. We included only studies that had analyzed gene expression as mRNA from oral tissues and compared with appropriate normal oral tissues. This was to ensure comparable gene expression and to remove potential bias through tissue specific gene expression. Studies that utilized non-human tissues, DNA methylation studies (platform difference), cytological smears were excluded. Methylation array studies were excluded. Only samples from currently untreated OLP were included. Data pertaining to dermal LP were excluded. When annotation of the data was difficult, the data was excluded.

After a thorough search and excluding datasets as specified above, two datasets for OLP (GSE52130 and GSE38616) comparing against normal oral mucosal tissues were selected for further analysis.^[38,39]^ A total of 14 OLP samples and 14 matched control tissues from oral mucosa were obtained and used for this study. The datasets were downloaded from the NCBI GEO database (https://www.ncbi.nlm.nih.gov/gds). The Matrix files were downloaded and processed. For each study we extracted the GEO accession number, platform, sample type, and gene expression data. The microarray chip identifiers were transformed to other suitable Gene IDs including Entrez Gene identifiers for downstream analysis. Datasets were merged after annotation with the Entrez Gene identifiers. A suitable identification class for each sample (as either OLP or Normal) were assigned, and further analysis was carried out with the web-based tool www.NetworkAnalyst.ca

Normalization by Log_2_ transformation with auto scaling to each dataset was performed. Each dataset was then visually inspected using PCA plots for gross deviation. The individual analysis of each dataset was carried out using the Benjamini–Hochberg’s False Discovery Rate (FDR) with cut-off p-values of <0.05. Batch effect between the different dataset were accounted with the ComBat batch effect method.^[40]^ For detecting the significantly differentially expressed genes (DEGs) between OLP cases and controls. The combining p-value method was employed to elicit more information through the integration of studies. Stouffer’s method was used to get a sensitive result.^[41]^ We set a discovery significant value of <0.05 to discover the most significant DEGs. From this pool of DEGs were downloaded and the genes of interest were studied.

The genes that were identified to be DE and also associated with efferocytosis process where subjected to functional enrichment analysis with literature keyword of “Efferocytosis” at GeneCllip3 website. In order to increase the sensitivity, filters of P≤1X10-10, hits≥5 was used. The clusters, enrichment score and the association were given as heat map. The subset of DE expressed genes associated with efferocytosis were subjected to literature based interaction analysis using Geneclip3 and interaction presented.

## RESULT

In the gene selection, from Genclip3, 243 genes were identified from 296 manuscripts that were associated with efferocytosis. From the www.genecards.org, 70 genes associated with efferocytosis were identified. From the genclip3, 243 genes were identified and from geneontology website, set of 132 genes were identified. From manuscripts, 128 genes were identified. After collating all the genes and removing duplicates, 304 genes related to efferocytosis remained.

Only 2 studies fulfilled the inclusion and exclusion criteria laid down.^[38,39]^ The individual patient level data were collected as outlined in the material section. In all there were 28 cases – 14 in oral lichen planus and 14 normal controls. For the meta-analysis, the cut-off p-values were adjusted using the Benjamini–Hochberg’s False Discovery Rate (FDR), in both studies and the same was fixed at 0.05.

In the first study, 47231 genes were annotated of which, on subject to DE analysis between OLP and normal, 2193 genes emerged significant while in the second study, there were 33297 genes annotated of which none of them emerged significant between OLP and normal. After combing the p-value, 1486 genes emerged DE between OLP and normal tissues.

The DE of genes associated with efferocytosis between OLP and normal are depicted in Table-2. Gene product of ADAM9 (P=0.048), ANXA2(P=0.048), APOE(P=0.012), NRP2 (P=0.027), PLAUR (P=0.05), S1PR5(P=0.018), TNFB3 (P=0.04), ABCA1 (p=0.018), IL1B2p=0.009), ITGAV(P=0.036) and PBX1(P=0.008) were statistically significant. The genes such as HMGB1 that had significance in one study lost, when the P-values were combined. All the mRNA of DE genes, their mean±SD, combined T-statistics and combined P-values are presented. (Table-2).

**Table-2:**
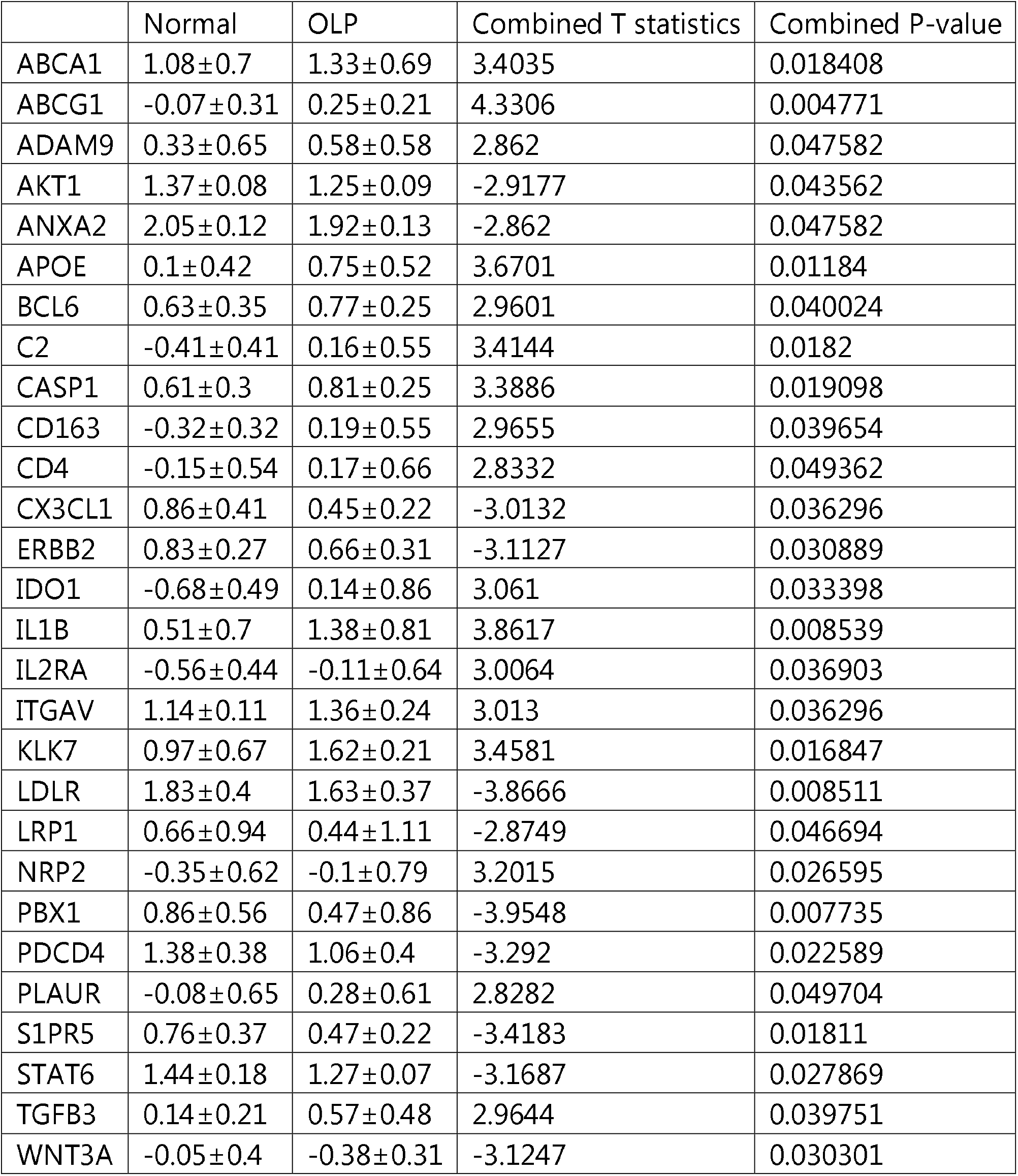
Log Transformed Expression of Significant mRNAs between Oral Lichen Planus (OLP) and Normal controls.

The functions of the DE mRNA genes were analyzed from an efferocytosis perspective using GenCLIP3. The same is depicted as a heat map in figure-2 (and data in supplemental file-1). There were 29 clusters identified (Supplemental file-1) with enrichment score ranging from 9.66 to 19.77. The network, protein-protein interaction of these DE mRNAs was also studied using literature curated GenCLIP3.(Figure-3). The strength of the association is reflected by the size of the nodes. The mRNA of AKT1, APOE, IL1B, PLAUR and LRP1 are more centrally placed than other genes while that of PBX1, ADAM9, NRP2 and KLK7 were placed at peripheries as they appear to interact with one or two mRNAs.

**Figure-2:**
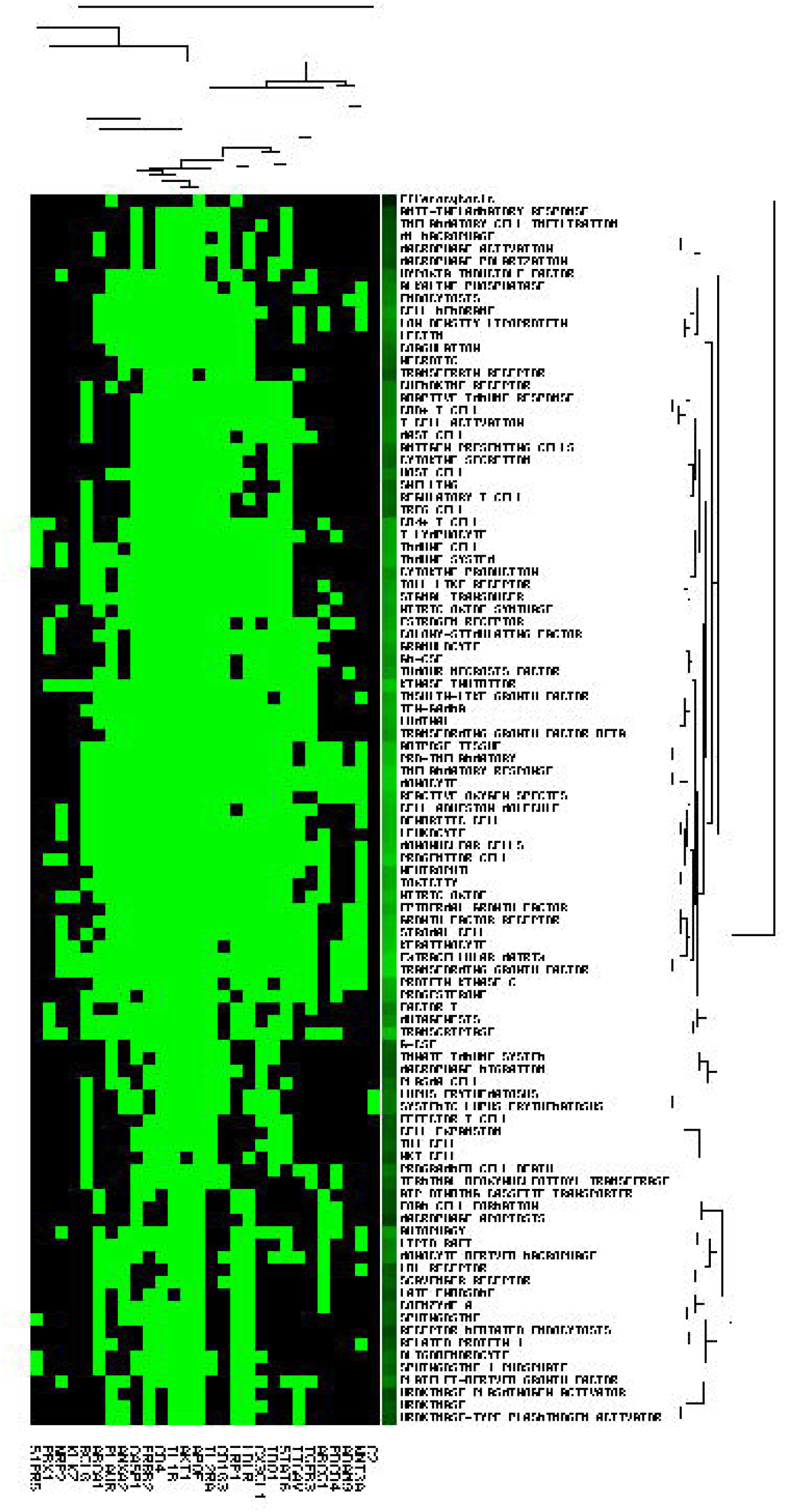
Heat map of differentially expressed gene’s mRNA and the various inflammation, apoptosis and cell death related functions

**Figure-3:**
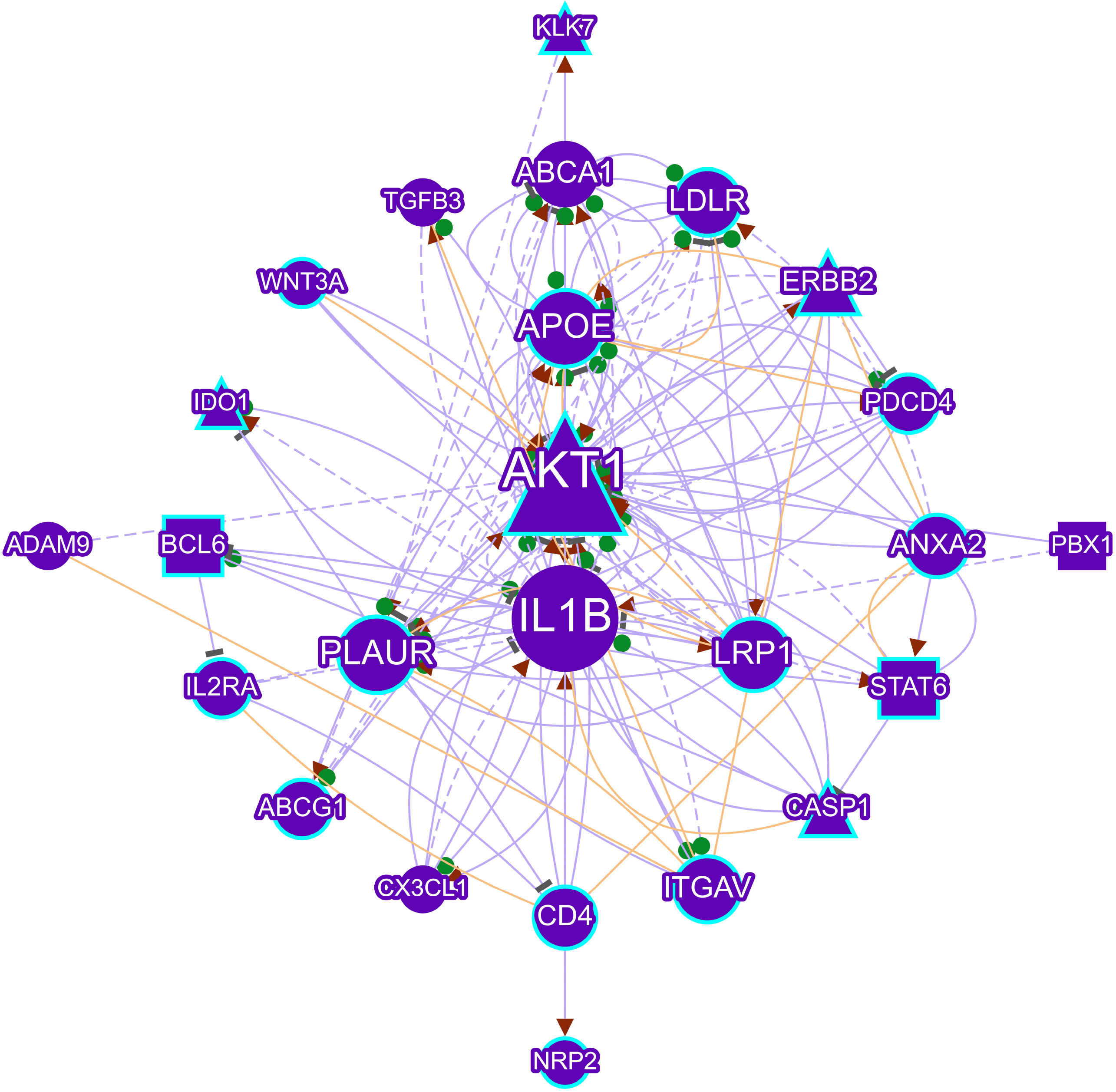
The relationship of differentially expressed genes with each other, as shown by GENCLIP3. [Violet, Round - Gene, Violet Triangle - Enzyme, Violet Square - Transcription factor, Violet Polygon - Enzyme and Transcription factor; Cyan blue outline - Genes associated with term, Green round and brown triangles - text mining, Brown - curated PPI database connection]

## DISCUSSION

Oral lichen planus (OLP) is an immune mediated, interface mucositis that exhibits hydropic degeneration of the basal cells, scattered dyskeratotic keratinocytes with a band like T lymphocyte rich inflammatory cell infiltrate near to the epithelial interface. The gene expressions altered in OLP has been previously reported.^[38,39]^ Even with such insights, the etiopathogenic mechanism of OLP still remains unclear.^[1-3]^ The death of keratinocytes is attributed to apoptosis and recently necroptosis.^[1-3,5]^ Recent studies have also postulated that pre-death stress determines the type of efferocytosis.^[9]^ This study was taken to identify the possibility of alterations in defective efferocytosis in OLP, as compared to normal.

For this purpose, we used mRNA analysis of tissue biopsy of OLP. Whole biopsy would contain oral keratinocytes and connective tissue with the rich inflammatory infiltrates. In contrast, smears would be devoid of crucial inflammatory cell infiltrates, which play a pivotal role in OLP. There are few studies that compared OLP with control, but suffered from several drawback including small sample size or considering whole genomic differences. For this study, we combined studies to have 14 OLP samples and equivalent controls using the latest statistical approach to identify combined P-values.

This study identified 1486 gene mRNA that were differentially expressed (DE) between OLP and controls. These could either be from the inflammatory cells or from the dying keratinocytes. The inflammatory cells, in a bid to suppress inflammation, would be signaled to initiate apoptosis and subsequent efferocytosis. In this process they would exhibit apoptotic and efferocytotic mRNA/proteins. Also, the dead and dying oral keratinocytes in OLP may express efferocytotic signals. Abnormal efferocytosis signals could further possibly delay inflammation resolution and may contribute to the chronicity of disease process.^[42,43]^

From the interaction perspective, AKT1 has a major role in efferocytosis via the unresolved endoplasmic reticulum stress and apoptosis. Macrophages that are involved with clearing of the cellular debris are resistant to apoptotic stimuli, because of activity of the PI3K/Akt pathway. Of the three types of AKT, AKT1 promotes M1-phenotype macrophage, which is pro-inflammatory and associated host defense. Reduction of AKT1, as in OLP, indicates reduced inflammation, indicating possibility of reduced efferocytosis and altered inflammatory signaling pathway.^[44-46]^

In OLP, reduction of ANXA2 and increase of C2 mRNA expression indicates reduction or failure in the recognition of ACs via the PS pathway.^[11-13]^ PBX1 is associated with IL10 expression and with immune tolerance in T cells.^[47]^ Reduced PBX1 in OLP could be indicative of altered immune tolerance and deranged T cell function. Altered PBX1 is known to predispose to systemic lupus erythematosus.^[48]^

Down-regulation of CX3CL1 in OLP, indicates the possible reduction of efferocytosis. ITGAV, through which the interginα5-intergrinβ3 binds to CX3CL1, is increased. As CX3CL1 is downregulated in OLP, signaling is dampened. In the second phase, the efferocyte receptors directly bind to AC (notably LRP1) and through bridging molecules (αvβ3 and αvβ5 integrins) along with ABCA1 and ABCG1.^[11]^ In OLP, LRP1 is reduced as compared to normal while ABCA1 and ABCG1 are increased. LRP1 being a pro-efferocytosis entity, reduction of LRP1 may reduce the efferocytosis as well as inhibit suppression of cell death. After cell death cholesterol is released, which needs to be removed by a process mediated by ABCA1 and ABCG1. In the LRP1 mediated, LXR–ABCA1 pathway, PPARG and its ligands, mediate cholesterol efflux.^[11]^ Increase of PLAUR, integrins and decreased LRP1 levels in OLP as compared to normal tissues indicate that the efferocytosis signals are there but possibly not effective or delayed.^[11]^

In OLP, APOE levels are increased but LRP1 levels are decreased, potentially altering the efferocytosis. Also, APOE would be increased when the inflammatory cells such as macrophages undergo apoptosis. Possibly, the increase of APOE could be from the dead or dying inflammatory cells. APOE binds to several receptors, such as LRP1 and LDLR. They can bind to ligands and can both internalize cargo and trigger signaling-mediated downstream effects, leading to efferocytosis. Reduction of LRP1 and LDLR in OLP will possibly impair efferocytosis.^[11,25]^

Increase in expression of IL1B, IL2RA and CASP1 possibly reflects AC as an ICD. ^[12,33]^ CD163, along with IL10, is a widely accepted marker of the M2-macrophage phenotype, that anti-inflammatory, immunosuppressive, and wound-healing. CD163 is increased in OLP.^[16,17]^ In OLP, as the AKT1 is decreased and CD163 is increased, possibly there is a shift of M2 macrophage phenotype. CD4 expressing T cell subset is known to increase efferocytosis by associating with IL10. This is increased M2-macrophage phenotype in OLP, indicates recruitment of CD4-T cells is a feature in a subset of OLP, as reported previously.^[17]^

Increased expression of ADAM9 and reduced STAT6 in OLP, would possibly lead to suppression of Mer-dependent phagocytosis, affecting intracellular downstream signaling, leading to defective/delayed efferocytosis and inflammation resolution.^[19,20]^ Wnt signals received by macrophages are important for their various phagocytic roles, such as modulating the immune response in the setting of infection, tissue repair after injury, malignancy detection, and progression. Wnt-3A mediates the Beta catenin signaling pathway.^[26]^ Reduction of this signaling mRNA expression, possibly indicate that the immunological response is altered.

During infection, a loss or reduction of PDCD4 protects from a bacterial enterotoxin. miR-21/PDCD4 axis modulates efferocytosis by the induction of anti-inflammatory “clean-up” genes such as interleukins.^[28,29]^ In OLP, this PDCD4 is reduced, indicating a possibility of a defective efferocytosis.

In ACs, S1PR5 blocks efferocytosis. S1PR5 of the circulating monocytes is possibly a by stander during tissue inflammation, exerting its influence on efferocytosis.^[31]^ Reduced S1RP5 in OLP could possibly relate to altered efferocytosis. KLK7 has a central role in desquamation and it catalyzes the degradation of intercellular cohesive structures. Also, it is associated with pro-inflammatory cytokine IL-1β, indicating its role in effective efferocytosis, as seen in increased levels in OLP.^[32]^ Tolerance to AC is regulated by IDO1, creating local immune suppression, maintains functional immune status in inflammatory scenarios.^[34]^ The increased mRNA expression of IDO1 in OLP could possibly be related to dysregulated efferocytosis.

In inflammatory diseases, promoting neutrophil apoptosis is one of the approach for the resolution of inflammation. ErbBs have known roles in suppressing apoptosis of epithelial cells and keratinocytes and ErbB2 reduction is known to resolve inflammation.^[35]^ In the present study we observed reduction of ErbB2, which is possibly suggestive of inflammation resolution. BCL6, mediates B cell differentiation, inflammation, inhibits differentiation and promotes proliferation.^[36]^ In OLP, possibly, the reduction of inflammation via the BCL6 is reflective of dysregulated efferocytosis.

Suppression of LRP1, LDLR, ANAX2, C2, PBX1, PDCD4, S1PR5, CX3CL1, STAT6 and Wnt3A indicates the possibility of defective or delayed efferocytosis in OLP. The interaction of the genes, indicating function and the pathways as reflected in the heat map strengthens our assumptions. The results of this study indicate that OLP has altered mRNA expression of certain genes/proteins that could potentially delay or lead to defective efferocytosis. Such a situation, trigger the inflammation in a waxing and waning pattern.

The result of the study needs further confirmation as the study is limited in its secondary data design, non-consideration of stage of OLP, type of OLP, and whole tissue approach. Single cell mRNA analysis of basal cells of OLP could possibly yield more insights.

## CONCLUSION

Whole OLP tissue mRNA expression were studied for possible alteration in defective or delayed efferocytosis. Alteration of crucial gene mRNA expressions were identified from this study. The results of this study need to be confirmed by efferocytosis specific markers, using a large scale sample at the epithelial, interface, connective tissue and inflammatory cell level to identify the role of the efferocytosis in the pathogenesis of OLP. This would help to design better treatment modalities. \

## Notes

### Competing Interest Statement

The authors have declared no competing interest.

